# Hi-C detects novel structural variants in HL-60 and HL-60/S4 cell lines

**DOI:** 10.1101/482208

**Authors:** Elsie C. Jacobson, Ralph S. Grand, Jo K. Perry, Mark H. Vickers, Ada L. Olins, Donald E. Olins, Justin M. O’Sullivan

## Abstract

Cancer cell lines often have large structural variants (SVs) that evolve over time. There are many reported differences in large scale SVs between HL-60 and HL-60/S4, two cell lines derived from the same acute myeloid leukemia sample. However, the stability and variability of inter- and intra-chromosomal structural variants between different sources of the same cell line is unknown. Here, we used Hi-C and RNA-seq to identify and compare large SVs in HL-60 and HL-60/S4 cell lines. Comparisons with previously published karyotypes identified novel SVs in both cell lines. Hi-C was used to characterize the known expansion centered on the MYC locus. The MYC expansion was integrated into known locations in HL-60/S4, and a novel location (chr4) in HL-60. The HL-60 cell line has more within-line structural variation than the HL-60/S4 derivative cell line. Collectively we demonstrate the usefulness of Hi-C and with RNA-seq data for the identification and characterization of SVs.

## 1 Background

Karyotyping with or without fluorescence *in situ* hybridization (FISH) are effective ways of identifying large scale structural variation, but both methods have relatively low resolution [1]. Genome wide chromatin conformation capture (Hi-C), which was developed to identify spatial genome organization [2,3], is emerging as a useful method for de-novo genome assembly [4] and identifying structural variants [5,6].

The HL-60 cell line was derived from a female acute myeloid leukemia patient in 1976 [7]. Since that time, the HL-60 cell line has been instrumental to breakthroughs in cancer, immunology, and cell biology. For example, HL-60 cells were used to identify retinoic acid as a therapeutic for acute myeloid leukemia [8,9], investigate myelocytic differentiation [10,11], study neutrophilic behaviors such as migration and respiratory bursting [12–15], and understand the mechanical properties of nuclear envelope components [16,17].

Several cell lines have been derived from the original HL-60 cells, including HL-T [18], HL-60/MX2 [19], and HL-60/S4 [20], however, the original HL-60 line [7] and the HL-60/S4 variant are currently used in empirical studies [20,21]. The HL-60/S4 subline was generated using random mutagenesis and selection for cells supersensitive to retinoic acid-induced differentiation [20]. A recent karyotype identified novel structural variants in the HL-60/S4 genome when compared to the parental HL-60 cell line [22].

A recent survey of MCF7 sublines showed rapid development of genetic diversity and subsequent functional changes in this breast cancer cell line, demonstrating that they cannot be used interchangeably for drug discovery screens or other purposes [23]. While HL-60 and HL-60/S4 are recognized as different cell lines by ATCC, they are both used for similar studies [22,24]. It is notable that despite the HL-60/S4 variant being developed as a super-responder to retinoic acid-induced differentiation; modern studies typically treat both HL-60 and HL-60/S4 with 1µM retinoic acid for four days to induce differentiation into the granulocytic form [22,24].

Cytogenetic information is available for both HL-60 and HL-60/S4 cell lines, however its utility is limited by two factors: 1) the resolution of SV detection by traditional karyotyping methods is low when compared to that achieved by sequencing-based methods [6]; and 2) genetic variation within cell lines [23] allows for the possibility that cell lines from different sources, or analyzed at different times, may not match the reference karyotype.

Here, we use sequencing data from multiple studies of two related cell lines to investigate SVs at high resolution. We validate the accuracy of our approach with published reference karyotypes, and investigate SVs that were newly identified in our study. Of these “novel” SVs, we identify two types: 1) small SVs that were likely present in the reference cell line, but remain undetected by traditional karyotyping methods; and 2) large SVs that are unlikely to have been missed by traditional karyotyping, and therefore may represent a divergence within the HL-60 cell line. The small SVs were predominantly found in HL-60/S4, and may represent updates to the current reference karyotype. By contrast the large SVs were both identified in the HL-60 Hi-C data. Therefore, in this manuscript we refer to the HL-60 datasets from Li et al 2018 [24] as describing HL-60/CICLR (HL-60 from the Chinese Infrastructure of Cell Line Resources). We investigate the composition and location of the known HL-60 MYC gene expansion. We confirm that it is highly stable in its composition, and its location in HL-60/S4 is consistent with the reference karyotype. However, we identify a novel integration of the MYC locus into chromosome 4 in HL-60/CICLR. Finally, we exploit the shape and orientation of SV patterns in Hi-C to characterize simple and complex SVs, two of which involve three or more chromosomes.

## 2. Results

### 2.1 Structural variants detected with Hi-C and RNA-seq

In order to investigate structural variation in HL-60/CICLR and HL-60/S4 we analyzed two types of sequencing data, Hi-C and RNA-seq, with structural variant (SV) detection tools. We then manually curated the outputs to identify a set of plausible SVs in each cell line.

Hi-C detects high frequency contacts between regions of DNA in close linear proximity, a feature which has been exploited to improve genome assemblies [4] and detect structural variation [5]. Very high contact frequencies between distal regions of the genome, therefore, likely represent a translocation or inversion compared to the reference genome [4–6].

We detected SVs from Hi-C data in three independent experiments across the two cell lines. Two independently generated Hi-C experiments were performed using HindIII and MboI on HL-60/S4 cells grown in RPMI 1640 with 1% pen/strep and 10% serum, in the same laboratory, two years apart [25]. An additional Hi-C data set, generated using HaeIII and HL-60/CICLR cells grown in RPMI 1640 with 1% penicillin/streptomycin and 10% serum [24] was retrieved from GEO GSE93997. The Hi-C data set from Jacobson *et al* [25] was generated using HL-60/S4 cells that had been differentiated into granulocytes using all-trans retinoic acid (ATRA) treatment for four days. Therefore, to avoid any treatment-specific effects, we only investigated the genomic structure in ATRA-treated HL-60 cells.

We used hic_breakfinder v1 [6] to identify structural variants from our Hi-C data. Regions of unusually high contact frequency, indicating a possible SV (‘breaks’), were identified in genome structures captured in the HL-60/CICLR [24], HL-60/S4-HindIII (this study) and in the HL-60/S4-MboI [25] datasets. To further investigate each of these breaks, we visually inspected the putative SVs in Hi-C contact matrices visualized in HiGlass [26]. We found that hic_breakfinder identified five short range intra-chromosomal breaks in the HL-60/S4-MboI data set that were not supported by visual inspection of the Hi-C heatmap. Therefore, we filtered the hic_breakfinder results to remove breakpoints that were less than 10Mb apart (Supp table 1). Of the 67 remaining breaks (Supp table 1), all except two (1—1, 3—8; both present in the HL-60/S4-MboI dataset [25]) were supported by visual heatmap inspection. In some instances, multiple break locations represent a single SV, resulting in a total of 27 identified SVs. Seven SVs were present in both HL-60/CICLR and HL-60/S4, two were unique to HL-60/CICLR, and 19 were unique to HL-60/S4 (Table 1).

**Table 1.** Locations of SVs identified within Hi-C datasets for HL-60-CICLR and HL-60/S4. Location 1 and 2 indicate the positions of the likely breakpoints. The next two columns indicate whether the SV was detected in HL-60/CICLR or HL-60/S4 respectively. A starred “yes*” indicates that this SV is not reported in the reference karyotype (DSMZ for HL-60/CICLR and Welch 2017 [22] for HL-60/S4). The Fusion Gene column indicates whether a fusion gene was identified at the SV. The Type column indicates the likely type of SV, based on Hi-C patterns and fusion transcripts where possible. The figure column indicates which figure the SV is shown in.

| Location 1 | Location 2 | HL60/CICLR | HL60/S4 | Figure | Fusion Gene | Strand | Distance | Predicted Type |
| --- | --- | --- | --- | --- | --- | --- | --- | --- |
| chr1 (p36.32) | chr3 (q28) | - | yes* | 1b |  |  | Inter | Translocation |
| chr2 (p24.2) | chr2 (p13.1) | - | yes* | 3b | SMC6--<br>SLC4A5 | -/- | Intra | Translocation |
| chr2 (p22.1) | chr2 (q31.2) | - | yes |  |  |  | Intra | Inversion |
| chr2 (p15) | chr2 (p13.1) | - | yes* | 3c |  |  | Intra | Inversion |
| chr2 (q14.1) | chr2 (q22.1) | - | yes* | S4a |  |  | Intra | Deletion |
| chr2 (q22.3) | chr2 (q24.3) | - | yes* | S4b |  |  | Intra | Inversion |
| chr3 (q28) | chr14 (q12) | - | yes |  | IL1RAP--AL136018.1 | +/+ | Inter | Translocation |
| chr4 (p15.31) | chr4 (p15.1) | - | yes* | 3c |  |  | Intra | Inversion |
| chr4 (p15.2) | chr8 (q24.13-q24.21) | yes* | - | 2b |  |  | Inter | Insertion |
| chr4 (q35.2) | chr18 (q21.1) | - | yes<sup>‡</sup> | 3c |  |  | Inter | Translocation |
| chr5 (q23.3) | chr5 (q31.3) | yes* | yes* | 3a |  |  | Intra | Deletion |
| chr5 (q31.2) | chr7 (q32.3) | yes | yes |  |  |  | Inter | Translocation |
| chr5 (q33.3) | chr16 (q23.2-q23.3) | yes | yes |  | CYFIP2--PLCG2 | +/+ | Inter | Translocation |
| chr5 (q11.2) | chr17 (p11.2) | yes | yes |  |  |  | Inter | Translocation |
| chr6 (p22.3) | chr6 (p12.1) | - | yes* | S4c |  |  | Intra | Inversion |
| chr6 (q22.2) | chr8 (q24.13-q24.21) | - | yes | 2b |  |  | Inter | Insertion |
| chr7 (q32.3) | chr16 (q24.1) | yes | yes |  |  |  | Inter | Translocation |
| chr8 (q24.13-q24.21) | chr11 (p14.1) | - | yes | 2b |  |  | Inter | Translocation |
| chr9 (q31.1) | chr14 (q23.2) | yes | yes |  |  |  | Inter | Translocation |
| chr10 (p12.1) | chr13 (q12.12) | yes* | - | 1a |  |  | Inter | Translocation |
| chr14 (q12) | chr21 (q22.11-q22.12) | - | yes* | 1c |  |  | Inter | Translocation |
| chr15 (q14-q21.1) | chr16 (p12.1-p11.2) | - | yes |  |  |  | Inter | Translocation |
| chr16 (p13.3-p13.11) | chr17 (q11.1-q25.2) | - | yes |  |  |  | Inter | Translocation |
| chr16 (p12.1) | chr21 (q22.11) | - | yes | 1c |  |  | Inter | Translocation |
| chr17 (q21.31) | chr17 (q25.3) | - | yes* | S4d |  |  | Intra | Inversion |
| chr18 (q21.1) | chr18 (q21.32) | - | yes | 3c |  |  | Intra | Deletion |
| chr20 (p11.23) | chr20 (q11.23) | - | yes* | S4e | RALGAPB--RBBP9 | +/- | Intra | Inversion |
| chr15 (q11.2) | chr21 (q22.11) | - | yes*<sup>†</sup> | 1c | TUBGCP5--TMEM50B | -/- | Inter | Translocation |

\*SV not reported in the reference karyotype (HL-60/CICLR reference from DSMZ, HL-60/S4 reference from Welch et al 2017)
<sup>†</sup>SV not detected by hic\_breakfinder in any dataset
<sup>‡</sup>Translocation reported in the reference karyotype, but not its reciprocal nature.

An alternative method to identify SVs is to look for gene fusions. To see if we could identify common and additional SVs from RNA-seq we used STAR-fusion v1.4.0 [27] on the HL-60/CICLR and HL-60/S4 cell transcription profiles. We analyzed HL-60/S4 gene fusions in two RNA-seq experiments from two different laboratories [22,28]. We analyzed HL-60/CICLR gene fusions in one RNA-seq experiment [24], the same source that used to acquire Hi-C data. All experimental datasets [22,24,28] included both undifferentiated and granulocytic (*i.e.* ATRA-differentiated) cells. STAR-fusion identified a total of 93 putative, unique gene fusions. To reduce the number of spurious fusion calls, we only considered gene fusions that were detected in all samples of a single condition, leaving 17 putative fusion candidates (Supp table 2). We then visually inspected the corresponding Hi-C heatmaps to identify SVs that could result in the predicted gene fusions. We identified one previously reported gene fusion in both HL-60/CICLR and HL-60/S4 [29], and four previously unreported gene fusions in HL-60/S4 (Table 1).

**Table 2.** Location (hg38) and gene content of the highly amplified regions surrounding MYC. These are consistent between Hi-C libraries, and with previous reports [37].

| Amplified regions within 8q24 |  |  |  |  |
| --- | --- | --- | --- | --- |
|  | Start (kb) | End (kb) | Coding genes | Non-coding genes |
| Amp 1 | 125,212 | 125,532 | NSMCE2,<br>TRIB1 | RN7SL329P;<br>AC084083.1;<br>AC091114.1 |
| Amp 2 | 125,700 | 126,372 |  | AC016074.1;<br>AC024681.2;<br>RNU6-442P;<br>KNOP1P5;<br>AC024681.1;<br>RFPL4AP5;<br>AC087667.1;<br>SOD1P3;<br>LINC00861 |
| Amp 3 | 127,056 | 127,331 |  | CASC19;<br>AC020688.1;<br>PRNCR1;<br>AC018714.1;<br>CASC8 |
| Amp 4 | 127,678 | 127,759 | MYC | CASC11 |
| Amp 5 | 128,989 | 129,202 |  |  |
| Amp 6 | 129,356 | 129,685 |  | MIR3686;<br>CCDC26;<br>AC103718.1;<br>AC011257.1 |

We found that hic_breakfinder had a low rate of false positive SV predictions above 10Mb in size, but that it was sensitive to Hi-C library preparation methods. In contrast, STAR-fusion had a relatively high false positive rate, although it was able to identify one likely SV that was not detected with hic_breakfinder. An integrative detection approach using both Hi-C and RNA-seq analysis tools is effective at identifying SVs, although manual assessment of each predicted SV is still required.

### 2.2 SVs reported in reference karyotypes are identified using Hi-C

In order to validate our approach, we compared our identified SVs to previously published karyotypes. We expect that even if some genetic variation exists within a cell line, most SVs will still be conserved.

Both HL-60 and HL-60/S4 cell lines have previously reported cytogenetic information (*e.g.* [22,30]), so we compared these to our Hi-C breakfinder results. The DSMZ catalogue describes the HL-60 karyotype (DSMZ no.: ACC 3), which we used as the reference for HL-60/CICLR. A recent study reported the karyotyping of HL-60/S4 using FISH [22], which is the karyotype reported in the American Tissue Culture Collection record of HL-60/S4 (ATCC^®^ CRL-3306^™^). All SVs described in the reference HL-60/S4 karyotype and all except two SVs (*i.e.* t(16;17), ins(1;8)) described in the reference HL-60 karyotype were identified with Hi-C, excluding duplications or whole chromosome deletion.

#### 2.2.1 Ancestral HL-60 SVs

As described in early HL-60 cell karyotypes [31], all HL-60 cells contained a set of SVs between chromosomes 5, 7, and 16 and a t(9;14) translocation. A translocation between chromosomes 5 and 17 was not reported in the spectral karyotype, but has been described elsewhere in an early HL-60 study [32] and is described in the DSMZ cytogenetic information. The SVs t(5;7)(q31.2;q32.3), t(5;16)(q33.3;q23.2-q23.3), t(7;16)(q32.3;q24.1), t(9;14)(q31.1;q23.2), and t(5;17)(q11.2;p11.2) were detected in both HL-60/CICLR and HL-60/S4 (Table 1). As these SVs are identified in all cytogenetic analyses of HL-60, to our knowledge, it is likely that they were present in the original leukemia sample.

#### 2.2.2 HL-60/S4 specific SVs

A recent karyotype using FISH reported additional SVs (*i.e.* inv(2)(p22.1;q31.2), t(3;14)(q28;q12), t(4;18)(q35.2;q21.1), t(16;21)(p12.1;q22.11)) specific to HL-60/S4 cells [22] that we identified in both Hi-C HL-60/S4-HindIII and HL-60/S4-MboI datasets. The previously reported t(15;16)(q14-q21.1;p12.1-11.2) and t(16;17)(p13.3-p13.11;q11.1-q25.2) translocations [22] were detected with hic_breakfinder in the HL-60/S4-HindIII dataset (this study) but not the HL-60/S4-MboI dataset [25] (Supp table 3). This apparent discrepancy may be due to restriction enzyme choice or other library preparation factors, especially given that both missing regions involve nearby regions of chromosome 16. For instance, MboI is methylation sensitive, while HindIII cuts at any site with the correct motif, regardless of methylation status. Thus, the sequencing coverage across breakpoints can be affected by the choice of restriction enzyme [33]. However, all SVs reported in the reference HL-60/S4 karyotype were identified with Hi-C, indicating both that the HL-60/S4 genome is stable and the detection tools are accurate.

**Table 3.** Summary of predicted gene fusions. The ‘SV’ column indicates whether there is evidence of an SV that could generate the gene fusion, based on Hi-C data.

| Fusion Gene | LeftChr | RightChr | Chromosomes | Distance | SV |
| --- | --- | --- | --- | --- | --- |
| AC098590.1--AC099789.1 | chr4 | chr1 | Different | >=10Mb | No |
| ACTG2--ACTG1 | chr2 | chr17 | Different | >=10Mb | No |
| AP004242.1--FOXP2 | chr11 | chr7 | Different | >=10Mb | No |
| CYFIP2--PLCG2 | chr5 | chr16 | Different | >=10Mb | Yes |
| IL1RAP--AL136018.1 | chr3 | chr14 | Different | >=10Mb | Yes |
| MTRNR2L12--AC025627.3 | chr3 | chr17 | Different | >=10Mb | No |
| PIBF1--KLF5 | chr13 | chr13 | Same | <10Mb | Unknown |
| RALGAPB--RBBP9 | chr20 | chr20 | Same | >=10Mb | Yes |
| RN7SKP71--RN7SKP48 | chr12 | chr4 | Different | >=10Mb | No |
| SLC7A5--SMG1P1 | chr16 | chr16 | Same | >=10Mb | Unlikely |
| SMC6--AC006030.1 | chr2 | chr2 | Same | >=10Mb | Yes |
| SMC6--SLC4A5 | chr2 | chr2 | Same | >=10Mb | Yes |
| SUZ12P1--TP53I13 | chr17 | chr17 | Same | <10Mb | Unknown |
| TMEM183A--PLEKHA6 | chr1 | chr1 | Same | <10Mb | Unknown |
| TUBGCP5--TMEM50B | chr15 | chr21 | Different | >=10Mb | Likely |
| USP22--RN7SL2 | chr17 | chr14 | Different | >=10Mb | No |
| USP34--ALMS1 | chr2 | chr2 | Same | >=10Mb | Unlikely |

### 2.3 Hi-C identifies non-canonical translocations in HL-60 and HL-60/S4 cell lines

Having verified that our approach was able to detect canonical translocations in HL-60/CICLR and HL-60/S4, we further investigated the translocations not described in the reference karyotypes.

In HL-60/CICLR, we detected a translocation between chromosomes 10 and 13 (t(10;13)(p12.1;q12.12), Fig 1a) that is not reported in the DSMZ reference karyotype. This translocation has been reported previously in HL-60 [34] but is absent from the HL-60 derived cell line HL-T [18]. Higher resolution inspection in HiGlass [26] confirms that this translocation is not present in HL-60/S4 (Supp fig 1a). The reference karyotype shows that HL-60/S4 cells have two complete versions of chromosome 10 [22], further indicates that this translocation was not present in the original leukemia sample. Moreover, the single comet tail indicates a non-reciprocal translocation, resulting in a derivative chromosome that consists of chromosome 10 (~25-133Mb) and chromosome 13 (~23-114Mb) (Fig 1a), a large ~200Mb chromosome that should be detectable with any cytogenetic analysis method.

**Figure 1.**
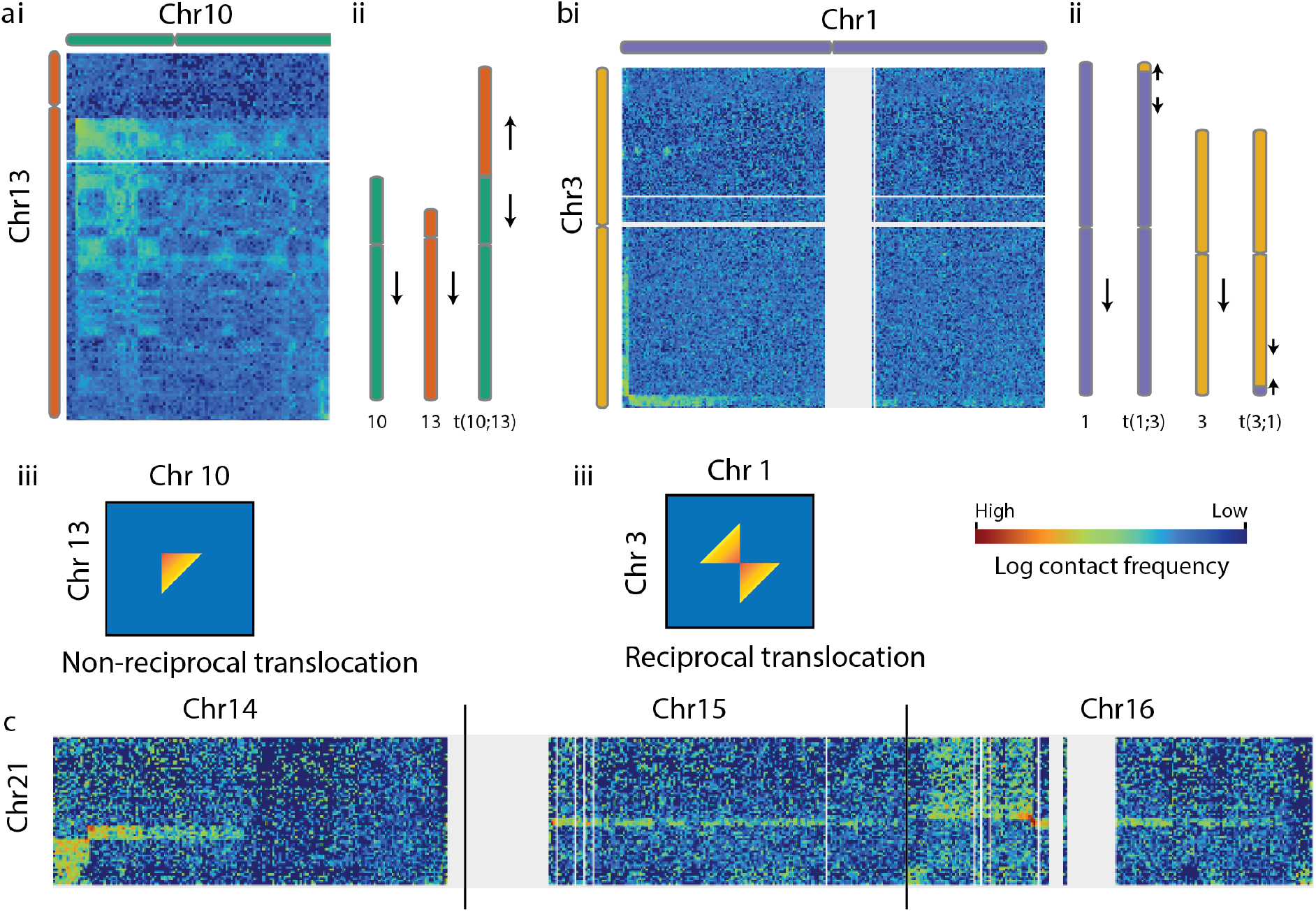
Non-canonical inter-chromosomal SVs in HL-60/CICLR and HL-60/S4. Hi-C heatmaps show the spatial contacts between two genomic locations, with the color of each pixel indicating the contact frequency. A) Characterization of t(10;13)(p12.1;q12.12) in HL-60/CICLR. i) The Hi-C heatmap of chromosome 10 and the q-arm of chromosome 13 show increased interaction frequency across the majority of both chromosomes, indicating a fusion of 13(q12.2-qter) onto 10(p12.1-qter). Ii) A single comet tail between two chromosomes indicates a non-reciprocal inter-chromosomal translocation. iii) Arrows indicated the reverse orientation of the translocated region. b) Characterization of t(1;3)(p36.32;q28) in HL-60/S4. i) Hi-C Heatmap of chromosomes 1 and 3 show increased interaction frequency between the start of chromosome 1 and the end of chromosome 3. ii) The pair of comet tails indicate a reciprocal translocation t(1;3)(p36.32;q28). iii) Arrows indicate the translocated regions have reversed orientation, as the p-arm of chromosome 1 has exchanged with the q-arm of chromosome 3. ci) Hi-C heatmap of the whole of chromosome 21 and chromosomes 14, 15, and 16, indicate a series of linked SVs in HL-60/S4. The previously unreported t(14;21)(q12;q22.11-q22.12) translocation has an unusual pattern: cii) Two comet tails are present, but one runs parallel to the diagonal, while the other runs perpendicular, indicating a complex translocation. ciii) The region immediately upstream of the t(15;21)(q11.2;q22.11) translocation is unmappable, making it difficult to interpret the heatmap pattern of this putative translocation. civ) The known t(16;21)(p12.1;q22.11) translocation [22] has a classical reciprocal translocation heatmap pattern, with the comet tails running parallel to the diagonal. Comparative heatmaps between cell lines are available in Supp fig 1.

In HL-60/S4, we identified previously unreported translocations between chromosomes 1 and 3 (t(1;3)(p36.32;q28), Fig 1b), and 14 and 21 (t(14;21)(q12;q22.11-q22.12), Fig 1c). Visual inspection of these putative SVs at multiple resolutions in HiGlass [26] confirmed that these translocations are not present in HL-60/CICLR (Supp fig 1b,c). While the HL-60 t(10;13)(p12.1;q12.12) translocation results in a large derivate chromosome, the t(1;3)(p36.32;q28) and t(14;21)(q12;q22.11-q22.12) variants observed in HL-60/S4 are much more subtle, and would require confirmation with breakpoint PCR or FISH. The paired comet tails indicate a reciprocal translocation between chromosomes 1 and 3. The chromosome 21q22.11 translocations with chromosomes 14q12 and 16p12.1 (Fig 1c) may have generated a derivative chromosome, or had two independent breaks on each homolog of chromosome 21. A further putative translocation was identified between chromosome 21q22.11 and chromosome 15q11.2. Despite hic_breakfinder not detecting a translocation between chromosomes 21 and 15 in either HL-60/S4-HindIII (this study) or HL-60/S4-MboI [25] datasets, there is a gene fusion (TUBGCP5--TMEM50B) identified across all HL-60/S4 samples and conditions that supports an SV between chromosomes 21q22.11 and 15q11.2. Heatmap inspection revealed poor sequence coverage at 15q11.2, which may have obscured the breakpoint from detection (Fig 1c). However, a ‘comet tail’ of increased chromosomal contacts can be seen to the right of the putative breakpoint, providing support for a translocation event.

The HL-60/CICLR t(10;13)(p12.1;q12.12) SV was not reported in the DSMZ cytogenetic information. Due to the large (~200Mb) derivative chromosome this translocation creates, its absence from cytogenetic information is unlikely to be due to usage of a low-resolution karyotyping method [35]. By contrast, the omission of the previously unreported inter-chromosomal SVs in HL-60/S4, which are small and/or complex, is likely due to the use of karyotyping methods that do not involve sequencing [6].

### 2.4 MYC expansion integrates into different locations within the genome

The *MYC* expansion that is present in HL-60 cells involves a non-contiguous region of chromosome 8 that is centered on the *MYC* gene (Fig 2) and amplified many times [32,36,37]. The region of chromosome 8q24 that comprises the *MYC* expansion has been reported to exist as: a) an extrachromosomal region; b) in the form of double minute chromosomes; and c) integrated into the genome [32,36–39]. We investigated both the composition and location of the MYC expansion using Hi-C and compared our results to previous characterizations of the locus.

**Figure 2.**
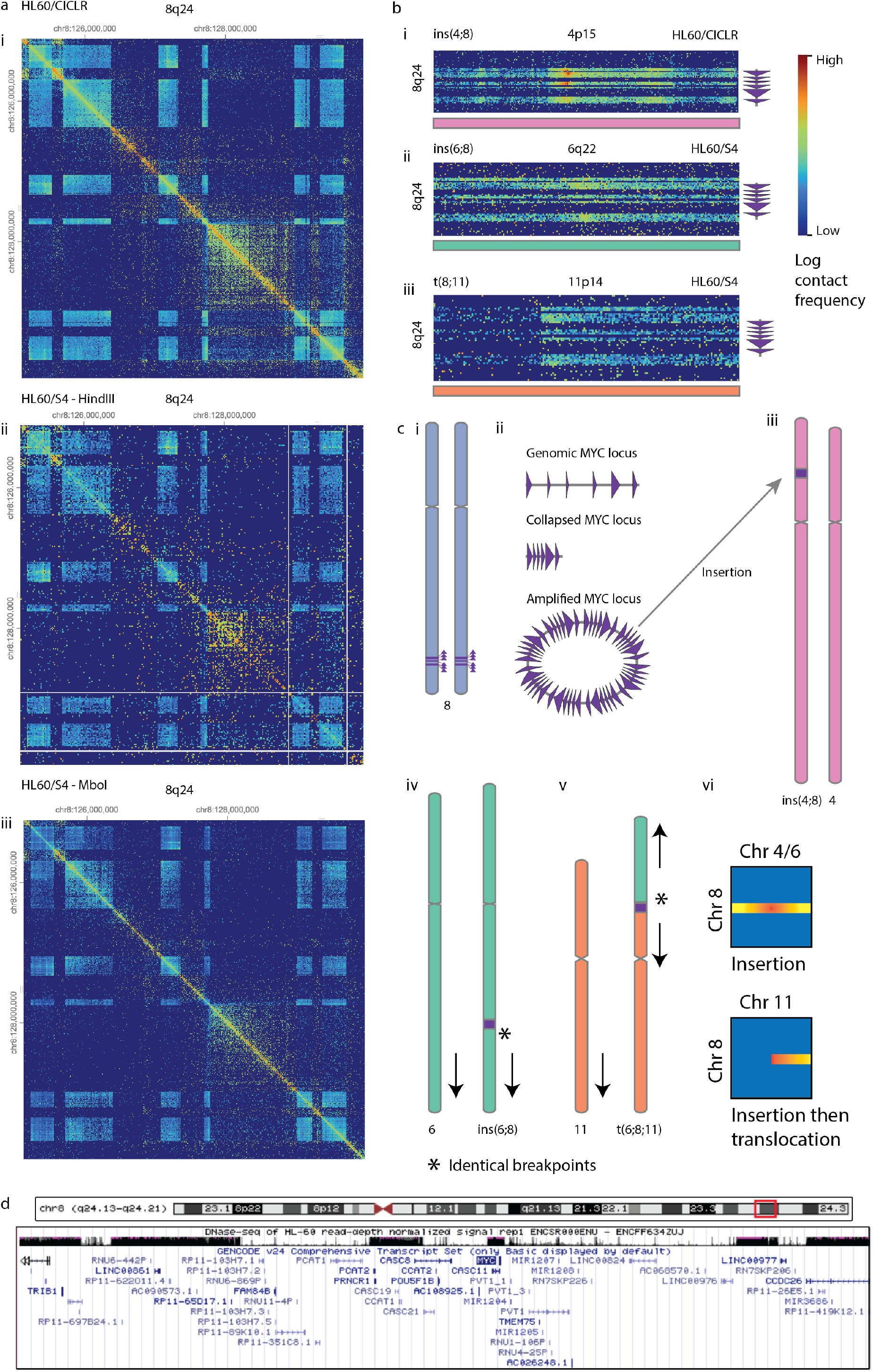
The MYC expansion and insertions at 8(q24.13-q24.21) were present in both HL-60/CICLR and HL-60/S4. a) Hi-C heatmaps have been ICE-normalized to adjust for variable coverage across different regions, but the massive copy number increase of the region centered on MYC has resulted in densely-population regions of the heatmap. The amplified regions are highly conserved across i) HL-60/CICLR, ii) HL-60/S4-HindIII, and iii) HL-60/S4-MboI. b) The amplified MYC expansion is i) inserted into chromosome 4 in HL-60/CICLR ii) inserted into chromosome 6 in HL-60/S4-HindIII and HL-60/S4-MboI, and iii) translocated onto chromosome 11 in HL-60/S4-HindIII and HL-60/S4-MboI. c) i) The MYC locus is a non-contiguous region of chromosome 8q24. ii) It lost the interleaving genomic regions, collapsing and amplifying as a circular extrachromosomal element, as described in Hirano 2008 [37]. iii) The amplified MYC locus has inserted into the p-arm of chromosome 4 in HL-60/CICLR, and (iv) inserted into the q-arm of chromosome 6 in HL-60/S4. v) The q-arm of chromosome 6, from the inserted MYC locus to the end of the chromosome, has then copied and translocated onto the p-arm of chromosome 11. Both ins(6;8) and t(6;8;11) can be seen in the karyotype of HL-60/S4 [22]. vi) The comet tail of the MYC insertion does not have the highest interaction frequency in one corner, but is distributed across the whole inserted locus. d) Genomic features of 8(q24.13-q24.21) (the MYC expansion). DNase hypersensitivity sequencing of HL-60 from ENCODE identifies the six amplified regions characterizing the MYC expansion. The MYC expansion is gene-dense, particularly with non-coding genes (Gencode v24 annotations). Comparative heatmaps between cell lines are available in Supp fig 2.

The composition of the *MYC* expansion is highly conserved between HL-60/CICLR and HL-60/S4 cells (Fig 2a), consistent with previous reports. The massive copy number increase means that in an ICE-normalized heatmap, amplified regions appear as densely populated ‘windows’ compared to the surrounding sparser matrix (Fig 2a). Due to the high density of these windows, the location of the six amplified regions that make up the *MYC* expansion can be identified at high resolution (Table 2). The presence of the six amplified regions is consistent with the amplicon described in [37], based on the size and genes contained in each amplified region (Table 2). Similarly, we observed high sequence coverage of the six amplified regions in normalized DNase hypersensitivity data from ENCODE (Fig 2d, [40]).

As reported in [22], the HL-60/S4 chromosome 8 is involved in two SVs, ins(6;8)(q22.2;q24.13-q24.21) and der(6;8;11)(q22.2;q24.13-q24.21;p14.1). Hi-C data of HL-60/S4 confirms that the *MYC* expansion is involved in translocations with chromosomes 6 and 11 (Fig 2b, Supp fig 2b). However, in the HL-60/CICLR cell line, the *MYC* expansion appears to be inserted into chromosome 4 (ins(4;8)(q35.2;q24.13-q24.21)) (Fig 2b, Supp fig 2a). One previous report has identified the *MYC* expansion integrating into t(5;17) in HL-60 [32]. By contrast, the DSMZ karyotype of HL-60 reports an insertion of 8q24 (containing the *MYC* expansion) into chromosome 1p31.

The difference between the insertions of 8q24 into chromosomes 4 and 6 and the translocation of 8q24 onto chromosome 11 is apparent in the Hi-C heatmaps (Fig 2b, [24,25]). Regions of increased interactions with 8q24 are observed upstream and downstream of the chromosomes 4 and 6 breakpoints. By contrast, there are no interactions upstream of the chromosome 11 breakpoint. Notably, there is only one breakpoint identified between chromosome 6 and chromosome 8. These results indicate that the derivative chromosome der(6;8;11) resulted from a translocation between chromosome 11 and the derivative chromosome ins(6;8), and not a translocation between chromosome 6 and 11 that was followed by an insertion of the *MYC* expansion (Fig 2c). Thus, the precise breakpoints identified with Hi-C can identify a likely timeline of genomic rearrangements in this instance.

Collectively, these indicate that the *MYC* expansion is highly conserved in its composition, but inserted into different regions of the genome in HL-60 derivative strains. There could be many reasons for this, but the simplest is that the extrachromosomal MYC expansion integrated into different regions of the genome in different clones of the original HL-60 sample, and has remained stably inserted during clonal expansion. However, there is some evidence to the contrary. Namely, an inversion identified in HL-60/S4 but not HL-60/CICLR inv(4)(p15.31;15.1)(Fig 3c) encompasses the location of the *MYC* expansion insertion (4p15.2), leaving open the possibility that the inversion is the result of *MYC* excision. If this is the case, the *MYC* expansion may continue to excise and insert into different genomic locations, and could even be responsible for other SVs identified in HL-60 genomes.

**Figure 3.**
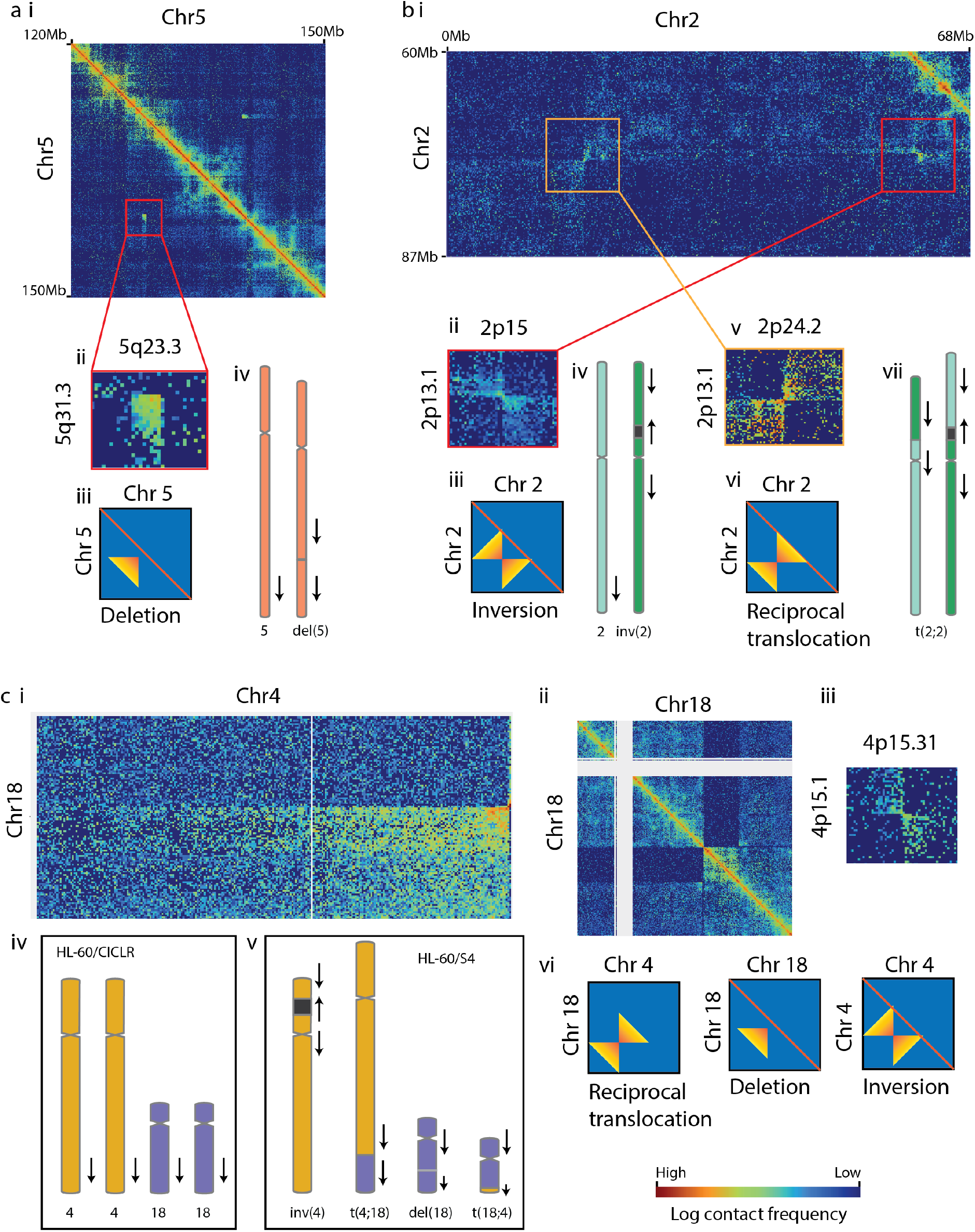
Non-canonical intra-chromosomal SVs in HL-60/CICLR and HL-60/S4. a) A deletion on chromosome 5 is absent from reference HL-60 karyotypes, but is present in both HL-60/CICLR and HL-60/S4. i) Hi-C heatmap of 30Mb region of chromosome 5, and (ii) is a zoom-in of the deletion. iii) The comet tail shows highest interaction frequency in the corner closest to the diagonal, is perpendicular to the diagonal, indicating a deletion. iv) The deletion is marked with a grey horizontal line in the q-arm of chromosome 5. It does not change the orientation of the chromosome, as indicated with arrows. b) Two apparently linked intrachromosomal SVs occur in chromosome 2 of HL60/S4. i) Non-symmetric Hi-C heatmap of a section of the chromosome 2 p-arm, with two SVs highlighted. ii) The 2(p13.1-p15) SV has a double comet tail, and the tails run parallel to the diagonal. iii) This is characteristic of an inversion. iv) The inversion reverses the orientation of 2(p13.1-p15), as indicated with arrows. v) The 2(p13.1-p24.2) SV has a double comet tail, but the tails run perpendicular to the diagonal, which is uncharacteristic of an inversion. Therefore, we propose the 2(p13.1-2p15) inverted region was reciprocally translocated onto the p-arm of the other chromosome 2 homolog, without changing chromosome orientation (vii). Additional intrachromosomal SVs on chromosome 2 are shown in Supp fig 4. c) Linked intra- and inter-chromosomal SVs on chromosome 4 and 18 in HL-60/S4. i) Hi-C heatmap showing a reciprocal translocation between chromosomes 4q35.2 and 18q21.1. ii) Hi-C heatmap showing a deletion on chromosome 18(q21.1-q21.32). There is a very low contact frequency between 18q21.1-q21.32 and the rest of chromosome 18. This is likely due to the t(4;18)(q35.2;q21.1) translocation removing the non-deleted allele from chromosome 18, resulting in interaction frequencies occurring at inter-chromosomal levels. iii) Hi-C heatmap showing an inversion on chromosome 4(p15.1-p15.3). Due to its distal location from the translocation, it is unlikely to be linked. However it is notable that the inverted region contains the MYC region in HL-60/CICLR (vi)(Fig 2). iv) The double comet tails in t(4;18) and 4(p15.1-p15.3) indicate a reciprocal translocation and inversion respectively, while the single comet tail trailing away from the diagonal indicate a deletion in chromosome 18. The translocation and deletion have comet tails perpendicular to the diagonal, indicating no change in orientation. The inversion comet tails run parallel to the diagonal, indicating a change in chromosomal orientation. v) Chromosomes 4 and 18 in HL-60/CICLR. The MYC insertion occurs at 4p15.2. vi) Chromosomes 4 and 18 in HL-60/S4, showing the 4(p15.1-p15.3) inversion, the 18(q21.1-q21.32) deletion, and the reciprocal t(4;18)(q35.2;q21.1) translocation.

### 2.5 Intrachromosomal SVs are sensitive to Hi-C detection

Inter-chromosomal SVs were generally identified across all samples with both karyotyping and Hi-C datasets. By contrast, intra-chromosomal SVs (*e.g.* inversions and deletions) were less likely to be reported in previously published karyotypes (*i.e.* [22]), and more likely to be missed by hic_breakfinder in one of the HL-60/S4 Hi-C datasets.

A deletion on chromosome 5(q23.3-31.1)(Fig 3a) was detected with hic_breakfinder in the HL-60-CICLR and HL-60/S4-MboI Hi-C datasets, and had weak visual evidence in the HL-60/S4-HindIII Hi-C dataset (Supp fig 3d). This deletion has been previously detected [31], but is not reported in either reference karyotype. However, its conservation between both HL-60/CICLR and HL-60/S4 provide support for it being an ancestral SV present in all modern strains. Notably, the chromosome 5 SV contains a cancer-associated locus (5q31.1) that is frequently deleted in non-small cell lung carcinoma [41].

Inversions on chromosomes 20(p11.23;q11.23) were identified with hic_breakfinder within the HL-60/S4-MboI Hi-C dataset, while an inversion on chromosome 17(q21.1;q21.32) was detected in HL-60/S4-HindIII Hi-C dataset (Supp fig 4d,e). A large pericentric inversion on chromosome 2(q22.1;q31.2) was reported in the HL-60/S4 reference karyotype [22], however, we detected four additional shorter-range SVs within chromosome 2 (*i.e.* t(2;2)(p24.2;p13.1), inv(2)(p15;p13.1) (Fig 3a, Supp fig 3a,b), del(2)(q14.1-q22.1), inv(2)(q22.3;q24.3) (Supp fig 4a,b))(Table 1). Three of these short-range SVs were identified by hic_breakfinder in the HL-60/S4-MboI Hi-C dataset, and one was identified in the HL-60/S4-HindIII Hi-C dataset. All four short-range SVs were visually confirmed in both HL-60/S4 Hi-C samples, but not in HL-60/CICLR (Supp fig 3a,b; Supp fig 4a,b).

Two of these SVs (inv(2)(p15;p13.1), inv(2)(q22.3;q24.3), Table 1, Fig 3c,e) have the distinctive pattern of reciprocal inversion, with paired comet tails running parallel to the diagonal. By contrast, the chromosome 2 SV (t(2;2)(p24.2;p13.1), Table 1, Fig 3b) has paired comet tails running perpendicular to the diagonal, consistent with it being a translocation between the two homologs of chromosome 2. A gene fusion arising from the t(2;2)(p24.2;p13.1) SV further indicates that the SV is the result of a translocation between homologous chromosomes, not an inversion. Specifically, gene fusions arising from inversions (e.g. *RALGAPB--RBBP9*) have different strandedness of the left and right parts of the chimeric transcript, as the region has been ‘flipped’. Both halves of the *SMC6--SLC4A5* fusion are negatively stranded, as is seen in fusions resulting from inter-chromosomal translocations such as *CYFIP2--PLCG2* (Table 1).

The inversion and translocation shown in figure 3b both involve 2p13.1, and are therefore likely to be linked SVs. One possibility is an inversion occurring in one chromosome, followed by DNA-damage induced recombination [42] between the two homologous chromosomes that would be undetectable without the inversion present.

A further complex set of SVs were identified within and between chromosomes 4 and 18 in HL-60/S4 (Fig 3c). The known translocation of 18(q21.1-qter) onto 4q35.2 was revealed to be a reciprocal translocation, with the terminal megabase of chromosome 4 translocated onto chromosome 18 (Fig 3ci). A large deletion in the same region of chromosome 18(q21.1;q21.32) was identified with hic_breakfinder in the HL-60/S4-MboI dataset. The deletion removed one allele of 18(q21.1;q21.32) entirely, and the translocation removed the other allele from chromosome 18, resulting in very low contact frequency between 18(q21.1;q21.32) and the rest of chromosome 18 (Fig 3cii). Finally, an inversion on chromosome 4(p15.1-15.3) (Fig 3ciii) is distal to the other SVs and therefore unlikely to be linked. However, it is notable that it encompasses the *MYC* insertion location in HL-60/CICLR (Fig 2bi, ciii, and 3c).

Intrachromosomal SVs are more challenging to identify than interchromosomal SVs. However, even with a detection limit of 10Mb, they constitute over a third of the SVs detected in this study. Moreover, the context (i.e. location and characteristics) of intrachromosomal SVs may help to shed light on the origins of some interchromosomal SVs.

### 2.6 Triaging putative gene fusions with Hi-C

Gene fusions are typically generated by genomic rearrangements [27]. Therefore, gene fusions detected from RNA-seq data should occur at SV breakpoints identified with Hi-C data. Of the 17 gene fusions identified across all replicates of at least one condition (Table 3), five unique predicted gene fusions occurred at breakpoints detected with hic_breakfinder, including two fusions located at the same breakpoint (SMC6--SLC4A5 and SMC6-- AC006030.1), and a sixth (TUBGCP5--TMEM50B) had visual support from Hi-C, despite poor sequence coverage across the region (Table 1, Fig 2cii). A further three predicted fusions had breakpoints less than 10Mb apart, and therefore their causal SV would not have been detected by our analysis. Two predicted fusions were between genes on the same chromosome, and so may have resulted from undetected SVs, given the reduced sensitivity of Hi-C to detect intra-chromosomal SVs. Six predicted fusions occurred between genes on different chromosomes, and had no corresponding SV detected with Hi-C data. Although it is possible that these gene fusions were generated by an alternative mechanism, such as trans-splicing [43], we predict these are false positives and would not prioritize these candidates for follow-up (Table 3).

## 3 Discussion

The aims of this study were threefold: Firstly, to identify structural variants and resulting gene fusions in two related cell lines (HL-60 and HL-60/S4). Secondly, to compare the SVs with existing karyotypes in order to assess the accuracy of this approach. Thirdly, to investigate genetic variation and evolution within HL-60 and its derivative cell lines.

In HL-60/CICLR, we found that most SVs were shared with the DSMZ reference, with notable exceptions. HL-60/CICLR contains a t(10;13) translocation, while the DSMZ reference contains a t(16;17) translocation. In HL-60/CICLR, the MYC expansion is inserted into chromosome 4, while the DSMZ reference reports it inserted into chromosome 1. While the functional importance of these differences cannot be known without further experiments, they are sufficiently large genomic alterations that HL-60/CICLR could be considered a distinct cell line from the DSMZ version of HL-60.

In HL-60/S4, we identified all SVs reported in the reference karyotype, as well as 10 previously unreported SVs, that were likely too small to be detected by FISH karyotype [22]. In contrast to HL-60/CICRL, we believe these novel SVs provide an updated view of the HL-60/S4 genome, as opposed to uncovering genetic diversity within the cell line. We found that detection of intra-chromosomal SVs was more sensitive to Hi-C method than inter-chromosomal SVs. We identified four novel gene fusions in HL-60/S4, including one that identified a likely translocation that was not identified with Hi-C.

We confirmed the conserved composition of the known *MYC* expansion, and have demonstrated the utility of Hi-C in identifying genomic insertion events. We also identified an inversion encompassing the HL-60/CICLR MYC location (4p15.31-p15.1) in HL-60/S4, providing possible evidence of *MYC* insertion and excision contributing to HL-60 genome evolution.

Hi-C data is predominantly generated with the purpose of investigating the spatial organization of genomes [2,3], but the same data can be repurposed to investigate genome structure without any additional wet-lab work [4–6,44,45]. The complexity of the genomic data that Hi-C generates has many uses and it represents a resource for SV detection.

Previous reports have identified the difference between reciprocal and non-reciprocal translocations based on Hi-C heatmap patterns [5]. In our analysis, we identified that four out of the eleven intra-chromosomal SVs in HL-60/S4 cells were not inversions (Table 4), based on the orientation of the butterfly block. In one instance, this was supported by the strandedness of a gene fusion that resulted from the SV (Table 1). In some instances, the complexity of the SVs, such as those seen with chromosome 21q22.11 and 18q21 in HL-60/S4, were difficult to interpret using Hi-C datasets alone. However, these SVs were also in some instances undetected by FISH karyotyping, presumably due to the small size of the rearranged regions. Analyzing Hi-C datasets was, therefore, able to highlight these for further, targeted analysis.

**Table 4.**
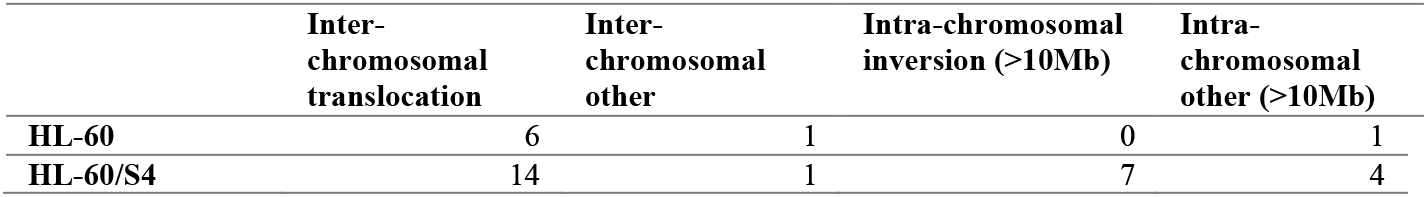
Summary of SVs in HL-60/CICLR and HL-60/S4 cells.

A recent study demonstrated that inter-chromosomal SVs could be detected precisely with Hi-C in a mouse erythroid cell line [46] and primary human brain tumors (five glioblastomas and one anaplastic astrocytoma). However, they did not report any intra-chromosomal SVs [5]. An SV detection tool developed by Dixon et al [6] was able to identify both intra- and inter-chromosomal structural variations. However, they did not use the Hi-C data to distinguish different forms of structural variation. Instead, they utilized optical mapping to characterize SVs. While this is preferable, patterns in Hi-C heatmaps can also be used to differentiate inversions from other forms of intra-chromosomal SVs without additional costs.

In this study we were able to characterize SVs based on the number and orientation of comet tails in Hi-C heatmaps. A pair of comet tails emanating from the same breakpoint indicates a reciprocal SV, while a single comet tail identifies a non-reciprocal SV. Comet tails running perpendicular to the diagonal identify SVs that do not change the orientation (*e.g.* deletion), while comet tails running parallel to the diagonal indicate a reversed orientation (*e.g.* inversion). Both orientations were seen in inter-chromosomal translocations. For example, the parallel comet tails in t(1;3)(p36.32;q28) indicate a reversed orientation, confirmed by the switching of the p and q arms. In contrast, t(4;18)(q35.2;q21.1) have comet tails running perpendicular to the diagonal, and indeed the translocation involved the q-arms of both chromosomes, supporting a translocation without a changed orientation. This may seem unnecessary information in the simple case, but the complex SVs with chromosome 21 highlight the importance of observing the shape and direction of comet tails.

Considering all aspects of the Hi-C, RNA-seq, and karyotype data allowed us to investigate complex SVs involving multiple rearrangement events. We were able to identify the likely order of the *MYC* locus traversing the HL-60/S4 genome, first inserting into chromosome 6 then translocating to chromosome 11, and possibly excising from chromosome 4. We identified a possible scenario of rearrangements within and between the homologs of chromosome 2, without performing haplotype phasing [47]. We located a complex linked set of SVs between four chromosomes (14, 15, 16, 21) that was partially missed with karyotyping due to the small regions involved, and includes a breakpoint in a region with poor sequence coverage.

The automated SV detection tool [6] is highly effective at identifying SVs. However, substantial additional characterization requires visual inspection of the putative SVs. New tools could be developed to include the orientation and symmetry of SVs and, thus, determine whether they resemble a reciprocal or non-reciprocal translocation, inversion, or other form of SV.

In theory, copy number variation can be detected with Hi-C [5]. However, the different restriction enzymes (*i.e.* HindIII, MboI, HaeIII) used in each Hi-C sample analyzed here complicated the analysis of genomic coverage. Despite this, the dramatic copy number increase of a region centered on the *MYC* gene (8q24) created a Hi-C heatmap pattern, even after matrix balancing to adjust for varied coverage (Fig 1a). The composition of the region (Table 2) was consistent across all three datasets, and with the literature [37]. The *MYC* expansion was inserted into chromosome 4p15.2 in HL-60, but was involved in SVs with chromosomes 6q22.2 and 11p14.1 in HL-60/S4. The precise breakpoints that we determined with Hi-C allowed us to deduce the probable order of these two SVs (ins(6;8)(q22.2;q24.13-q24.21) and der(6;8;11)(q22.2;q24.13-q24.21;p14.1), Table 1, Fig 1c). While the location of the *MYC* expansion was stable across two HL-60/S4 samples, it does not appear to be stable across different HL-60 sources, with our analysis showing an insertion into chromosome 4p15.2, while the DSMZ cytogenetic information describes an 8q24 insertion into chromosome 1, a 1990 study found it inserted into a t(5;17) translocation, and a spectral karyotype reported an insertion of chromosome 8 into the p arm of chromosome 11, but no chromosome 6 insertion [31].

We propose that the *MYC* expansion either; a) was stably inserted into different genomic locations during seed-lot generation at different repositories, or b) is a mobile element that may be inserted and excised, contributing to HL-60 genome evolution. If the former is true, different sources of HL-60 should confirm and report the current location of the *MYC* expansion, and derivative strains named if their *MYC* expansion location is different from the canonical strain (*i.e.* HL-60/DSMZ). Further experiments should be performed to ascertain whether the *MYC* expansion remains able to move around the genome. For instance, HL-60 could be grown for many passages, possibly with the addition of a mutagenic agent to encourage homologous recombination. Samples would be collected at many time points and PCR performed with two forward primers aligned to the end of the *MYC* expansion, with random hexamers as reverse primers. This would amplify sequences that occur at a junction with the *MYC* expansion that can be sequenced to identify the site of insertion. This would confirm whether the *MYC* expansion is stably inserted, or whether it is mobile and warrants further investigation.

The *MYC* extra-chromosomal element is particularly well described in HL-60 cells, but *MYC* (and *MYC* homolog) extra-chromosomal amplifications have been identified in a number of different cancers, most commonly leukemias and glioblastomas [48–55]. *MYC* deregulation in cancer generally increases the expression levels of the *MYC* gene, maintaining cancer cells in a highly proliferative state [56–59]. Intriguingly, overexpression of MYC is sufficient to drive genomic instability and induce the formation of extra-chromosomal elements such as double minutes [56–60], meaning that amplification of *MYC* can drive further amplification of *MYC* in a positive feedback loop. If these amplifications can also contribute to genome rearrangements via insertion and excision of extra-chromosomal copies, this could provide a second mechanism by which *MYC* amplifications can contribute to genomic instability, and subsequently tumor development and progression.

Structural variants can have functional consequences in a number of ways. For example, the generation of gene fusions can affect the normal functions of the genes involved in the fusion, in some cases driving cancer progression (*e.g.* PML-RAR [61] and BCR-ABL1 [62]). SVs can also disrupt the local regulatory environment of a gene, exposing it to repressive or enhancer elements that alter its normal gene expression level (*e.g.* [63,64]). The distinct SVs in HL-60/CICLR, HL-60/DSMZ, and HL-60/S4 may have profound effects on cell behavior and responses. However, it is also likely that these cell lines have single nucleotide differences as well as large SVs [23], and distinguishing the phenotypic effects of a single mutation, large or small, would require further experiments. For instance, associating allele specific expression with the SV containing homolog would require genome phasing combined with RNA-seq variant analysis, and fusion protein analysis would require ectopic expression in a different cell line. Therefore, we have avoided speculating on the functional consequences of SVs identified in this study.

Hi-C alone is not sufficient to perform a comprehensive analysis of all structural variations within a cell line. Rather, comprehensive characterization of SVs requires validation and investigation from orthogonal techniques [6]. Despite this, Hi-C can provide substantial information regarding the location and type of SVs. Moreover, Hi-C and RNA-seq libraries are often generated together in gene regulation studies (*e.g.* [25,65,66]). As such, integrating automated detection tools, *e.g.* hic_breakfinder [6] and STAR-Fusion [27], with visual heatmap inspection can provide substantial information regarding genomic SVs without any additional cost.

## 4 Conclusion

HL-60/S4 cells have a stable karyotype, with a number of small SVs that are not detected with FISH-karyotyping. HL-60 cells from the China Infrastructure of Cell Line Resources are distinct from HL-60 cells from the DSMZ repository. The *MYC* expansion has a stable composition, but is found in a different genomic location in each HL-60 cytogenetic analysis, indicating it may act as a mobile element. Hic_breakfinder was effective at identifying inter-chromosomal translocations and intra-chromosomal SVs, including linked and complex SVs. Visual inspection of SV patterns in the Hi-C heatmap distinguished inversions from other forms of intra-chromosomal SV, which was supported by the gene fusions identified with STAR-Fusion. Hi-C and RNA-seq are effective tools to characterize the location and type of genomic SV.

## 5 Methods

### 5.1 Cell culture – HL-60/S4-HindIII

HL-60/S4 cells were maintained in RPMI 1640 medium, plus 10% fetal calf serum and 1% penicillin, streptomycin, and l-glutamine at 37°C in a humid incubator purged with 5% CO_2_/95% air. Differentiation of HL-60/S4 cells into granulocytes was induced by all-trans retinoic acid (ATRA) treatment for four days [25]. Briefly, cells were seeded at a density of 1×10^5^ cells/ml in the RPMI medium (above) and ATRA (1µM final concentration).

### 5.2 Hi-C – HL-60/S4-HindIII

Dilution Hi-C library preparation was adapted from Lieberman-Aiden *et al.* 2009 [2]. Briefly, 15×10^7^ cells were crosslinked with Formaldehyde (2% FC, 10 min, RT), quenched with Glycine (125mM, 5 min, RT) and scraped from the plate. Cells were lysed, digested (HindIII) and ligated according to Lieberman-Aiden *et al.* 2009 [2] (Supplementary methods). A single replicate was sequenced (Illumina Hi-Seq 2000, DKFZ, Heidelberg, Germany). Hi-C QC information is available in supplementary table 4.

### 5.3 Hi-C data processing

Reads were aligned to hg38 and filtered using HiCUP v0.5.9 [67]. SVs were detected with hic_breakfinder [6]. SVs were filtered and summarized in R [68] with dplyr [69]. Iterative correction and eigenvector decomposition (ICE) was performed with cooler v0.7.1 [70]. Hi-C heatmaps were generated with HiGlass v1.1.5 [26].

### 5.4 RNA-seq analysis

Reads were aligned and fusions detected with STAR-Fusion [27,71] with hg38 and gencode annotations v27. Fusions were filtered and summarized in R [68] with dplyr [69].

## 7 Additional files

Supplementary Figure 1. Comparative heatmaps of SVs shown in figure 1, showing the same region in HL-60/CICLR, HL-60/S4-HindIII, and HL-60/S4-MboI. a) The non-reciprocal translocation t(10;13)(p12.1;q12.12) is present only in HL-60/CICLR. b) The reciprocal translocation t(1)(p36.32;q28) is present only in HL-60/S4. c) The t(14;21(q12;q11) translocation is present in only HL-60/S4. Although the two comet tails indicate a reciprocal translocation, the tails do not trail off opposite each other, suggesting a more complex SV.

Supplementary Figure 2. Comparative heatmaps of SVs shown in figure 2, showing the *MYC* expansion (8(q24.13-q24.21)) associated SVs in HL-60/CICLR, HL-60/S4-HindIII, and HL-60/S4-MboI. a) The *MYC* expansion was inserted into 4p15 in HL-60/CICLR. There was no evidence of insertion at this locus in HL-60/S4. b) The *MYC* expansion was inserted into chromosome 6q22 in HL-60/S4, but not in HL-60/CICLR. c) The *MYC* insertion on chromosome 6 was translocated onto chromosome 11p14 in HL-60/S4.

Supplementary Figure 3. Comparative heatmaps of SVs shown in figure 3, showing the same region in HL-60/CICLR, HL-60/S4-HindIII, and HL-60/S4-MboI. a) A putative inter-homolog translocation at 2(p13.1-p24.2) is only present in HL-60/S4. b) The inversion at 2(p13.1-p15) is only present in HL-60/S4. c) The inversion at 4(p15.1-p15.31) is only present in HL-60/S4.d) The deletion at 5(q23.3-q31.3) is present in both HL-60/S4 and HL-60/CICLR. e) The deletion at 18(q21.1-q21.32) is only present in HL-60/S4.

Supplementary Figure 4. Heatmaps of previously unreported intra-chromosomal SVs in HL-60/S4 cells. a) The 2(q14.1-q22.1) deletion is present in HL-60/S4 but not HL-60/CICLR. b) The 2(q22.3-q24.3) inversion is present in HL-60/S4 but not HL-60/CICLR. c) An inversion on chromosome 6 is present in HL-60/S4, but not HL-60/CICLR cells. d) An inversion on chromosome 20 is present in HL-60/S4, but not HL-60/CICLR cells. e) An inversion on chromosome 20 is present in HL-60/S4, but not HL-60/CICLR cells.

Supplementary Figure 5. Genomic positions of SVs in HL-60 cell lines. a) HL-60/CICLR ICE-normalized heatmap and SV locations b) HL-60/S4 ICE-normalized heatmap and SV locations. The unclassified intrachromosomal SV on chromosome 18 (Supp fig 1b) has been grouped with the inversions in this figure.

Supplementary Table 1. Hic-breakfinder filtered results for all samples. This table shows all breakpoints over 10Mb apart. The log-odds score indicates the strength of the call. The bias value predicts which coordinate is closest to the breakpoint. A “+” bias indicates the “end” coordinate is closest to the breakpoint, while a “-” bias indicates the “start” coordinate is closest to the breakpoint. The resolution indicates the maximum resolution the breakpoint was identified at, as hic_breakfinder identifies breakpoints at multiple resolutions. The sample column indicates which dataset the breakpoint was identified in.

Supplementary Table 2. STAR-Fusion filtered results for all samples. This table has the gene fusion predictions for all samples of the 17 fusions identified in all replicates of at least one condition. Each row provides the STAR-Fusion results for a single RNA-seq library, thus each gene fusion may fill up to 29 rows. The first three columns indicate the dataset, condition, and replicate of the library. The remaining columns include detailed information about the predicted fusion as outputted by STAR-Fusion, including specific breakpoint locations and the number of junction and spanning reads.

Supplementary Table 3. Summarized locations of SVs identified within Hi-C datasets for HL-60-CICLR [24], HL-60/S4-HindIII (this study), and HL-60/S4-MboI [25] cell lines. Location 1 and 2 indicate the positions of the likely breakpoints. The next three columns indicate the highest level of evidence provided for the SV occurring in that sample, in the order: Hic_breakfinder (breakfinder), STAR-Fusion, visual inspection in HiGlass (visual), or none (-). The Fusion Gene column indicates whether a fusion gene was identified at the SV. The Type column indicates the likely type of SV, based on Hi-C patterns and fusion transcripts where possible.

Supplementary Table 4. HiCUP QC results for HL-60/S4-HindIII.

Supplementary methods. Further details of the HL-60/S4-HindIII Hi-C library preparation.

## 8 Declarations

### 8.1 Data availability

The HL-60/S4-HindIII Hi-C raw and processed data is available at GEO, GSE120815. The HL-60/S4-MboI Hi-C from [25] is publicly available on GEO, GSE115634. The HL-60/S4-mRNA from [22] is publicly available in the NCBI Short Read Archive at http://www.ncbi.nlm.nih.gov/bioproject/303179. The HL-60-CICLR Hi-C and HL-60 RNA from [24] is publicly available on GEO, GSE93997. The HL-60/S4-total RNA is publicly available on GEO, GSE120579.

### 8.2 Ethics approval and consent to participate

Not Applicable

### 8.3 Consent for publication

Not Applicable

### 8.6 Authors’ Contributions

ALO and DEO grew HL-60/S4 cells and RSG prepared the HindIII Hi-C library, under the guidance of JMO. ECJ analyzed and interpreted the data, and wrote the manuscript with support from JMO, JKP, and MHV. All authors contributed to revisions and approved the final manuscript.

## 8.7 Acknowledgements

The authors would like to thank anonymous reviewer three for their assistance in understanding the structural variation in chromosome 18.

