## Supplemental table 3 for "Hi-C detects novel structural variants in HL-60 and HL-60/S4 cell lines"

| Location 1 | Location 2 | HL60-HaeIII | HL60/S4-HindIII | HL60/S4-MboI | Fusion Gene | Strand | Distance | Predicted Type |
| --- | --- | --- | --- | --- | --- | --- | --- | --- |
| chr1 (p36.32) | chr3 (q28) | - | breakfinder | breakfinder |  |  | Inter | Translocation |
| chr2 (p24.2) | chr2 (p13.1) | - | STAR-Fusion | breakfinder | SMC6--SLC4A5 | -/- | Intra | Translocation |
| chr2 (p22.1) | chr2 (q31.2) | - | breakfinder | breakfinder |  |  | Intra | Inversion |
| chr2 (p15) | chr2 (p13.1) | - | breakfinder | visual |  |  | Intra | Inversion |
| chr2 (q14.1) | chr2 (q22.1) | - | visual | breakfinder |  |  | Intra | Deletion |
| chr2 (q22.3) | chr2 (q24.3) | - | visual | breakfinder |  |  | Intra | Inversion |
| chr3 (q28) | chr14 (q12) | - | breakfinder | breakfinder | IL1RAP--AL136018.1 | +/+ | Inter | Translocation |
| chr4 (p15.31) | chr4 (p15.1) | - | visual | breakfinder |  |  | Intra | Inversion |
| chr4 (p15.2) | chr8 (q24.13-q24.21) | breakfinder | - | - |  |  | Inter | Insertion |
| chr4 (q35.2) | chr18 (q21.1) | - | breakfinder | breakfinder |  |  | Inter | Translocation |
| chr5 (q23.3) | chr5 (q31.3) | breakfinder | visual | breakfinder |  |  | Intra | Deletion |
| chr5 (q31.2) | chr7 (q32.3) | breakfinder | breakfinder | breakfinder |  |  | Inter | Translocation |
| chr5 (q33.3) | chr16 (q23.2-q23.3) | breakfinder | breakfinder | breakfinder | CYFIP2--PLCG2 | +/+ | Inter | Translocation |
| chr5 (q11.2) | chr17 (p11.2) | breakfinder | breakfinder | breakfinder |  |  | Inter | Translocation |
| chr6 (p22.3) | chr6 (p12.1) | - | visual | breakfinder |  |  | Intra | Inversion |
| chr6 (q22.2) | chr8 (q24.13-q24.21) | - | breakfinder | breakfinder |  |  | Inter | Insertion |
| chr7 (q32.3) | chr16 (q24.1) | breakfinder | breakfinder | breakfinder |  |  | Inter | Translocation |
| chr8 (q24.13-q24.21) | chr11 (p14.1) | - | breakfinder | breakfinder |  |  | Inter | Translocation |
| chr9 (q31.1) | chr14 (q23.2) | breakfinder | breakfinder | breakfinder |  |  | Inter | Translocation |
| chr10 (p12.1) | chr13 (q12.12) | breakfinder | - | - |  |  | Inter | Translocation |
| chr14 (q12) | chr21 (q22.11-q22.12) | - | breakfinder | breakfinder |  |  | Inter | Translocation |
| chr15 (q14-q21.1) | chr16 (p12.1-p11.2) | - | breakfinder | visual |  |  | Inter | Translocation |
| chr16 (p13.3-p13.11) | chr17 (q11.1-q25.2) | - | breakfinder | visual |  |  | Inter | Translocation |
| chr16 (p12.1) | chr21 (q22.11) | - | breakfinder | breakfinder |  |  | Inter | Translocation |
| chr17 (q21.31) | chr17 (q25.3) | - | breakfinder | visual |  |  | Intra | Inversion |
| chr18 (q21.1) | chr18 (q21.32) | - | visual | breakfinder |  |  | Intra | Unclassified |
| chr20 (p11.23) | chr20 (q11.23) | - | visual | breakfinder | RALGAPB--RBBP9 | +/- | Intra | Inversion |
| chr15 (q11.2) | chr21 (q22.11) | - | STAR-Fusion | STAR-Fusion | TUBGCP5--TMEM50B | -/- | Inter | Translocation |
