## Supplementary figures and images for "Hi-C detects novel structural variants in HL-60 and HL-60/S4 cell lines"

### Supplemental figure 1

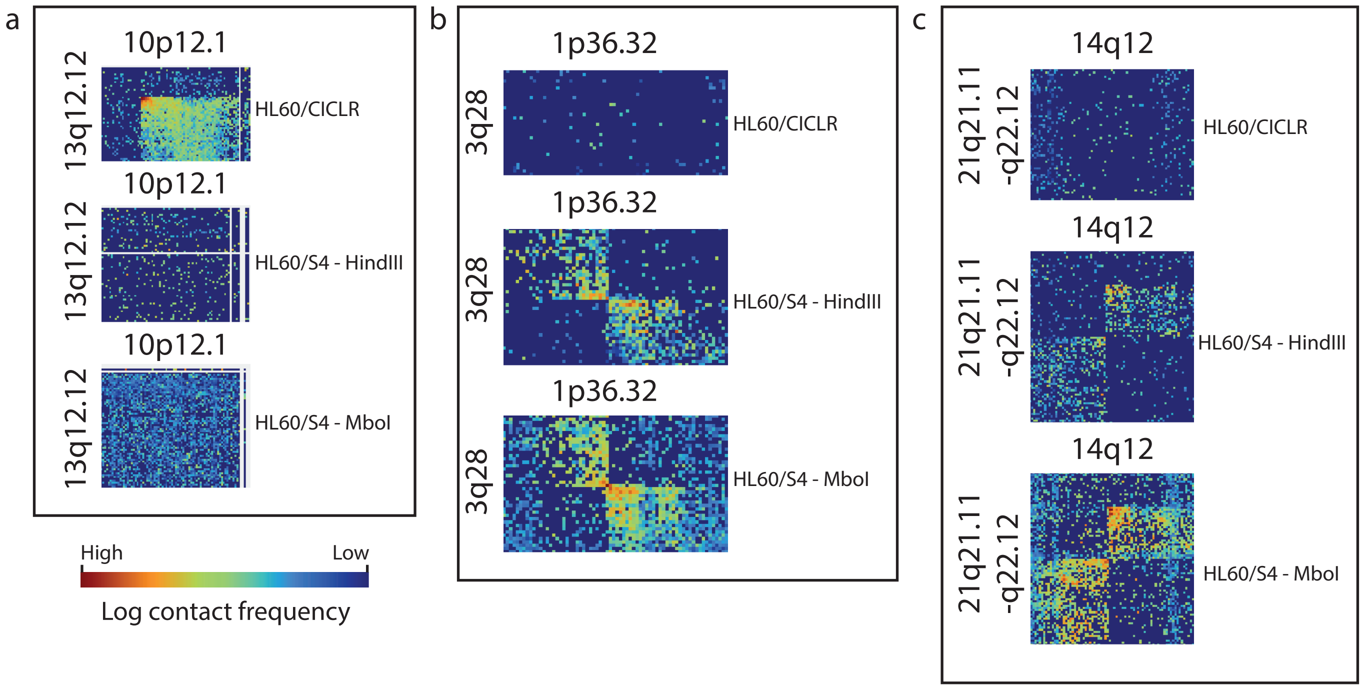

### Supplemental figure 2

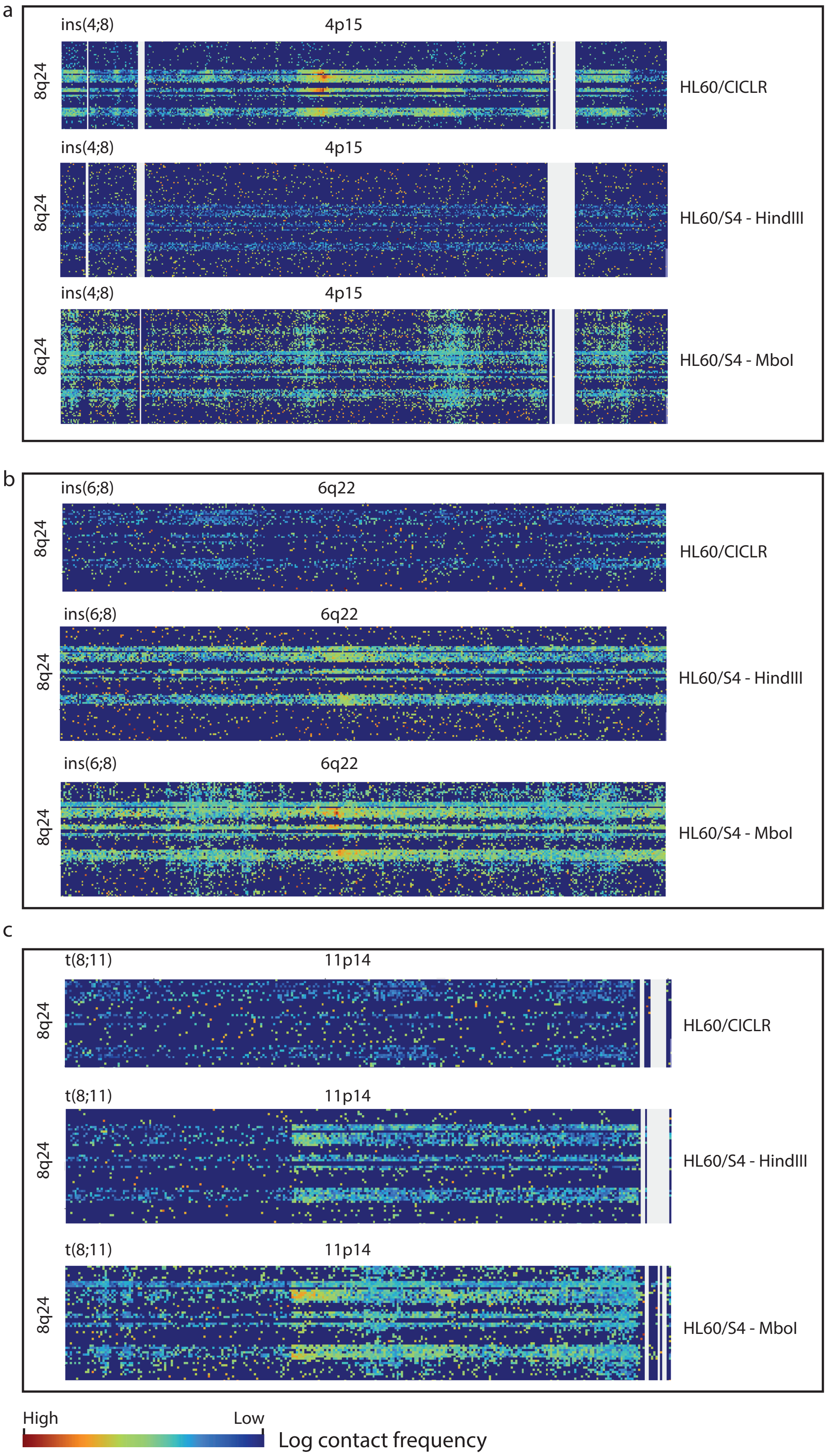

### Supplemental figure 3

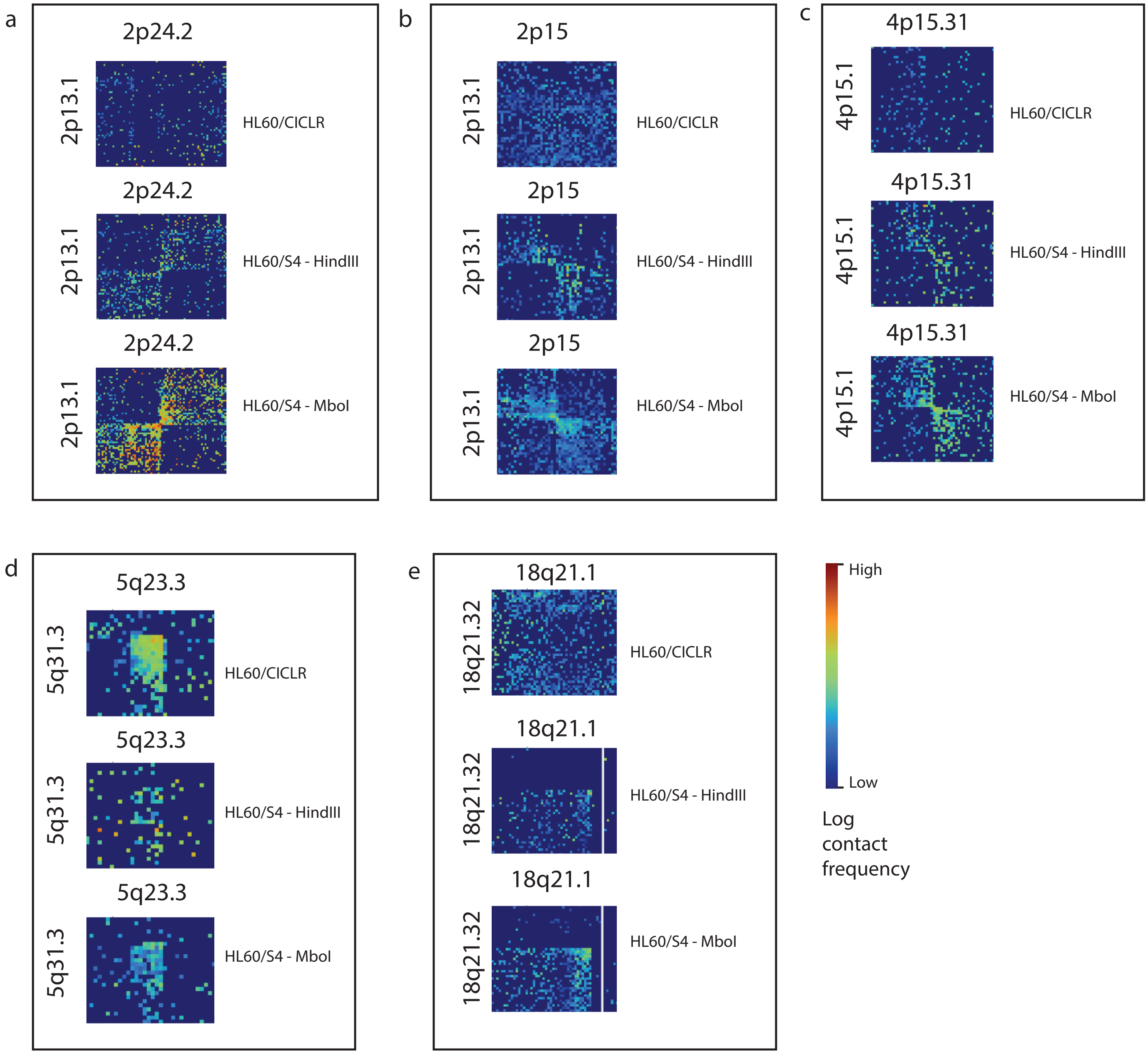

### Supplemental figure 4

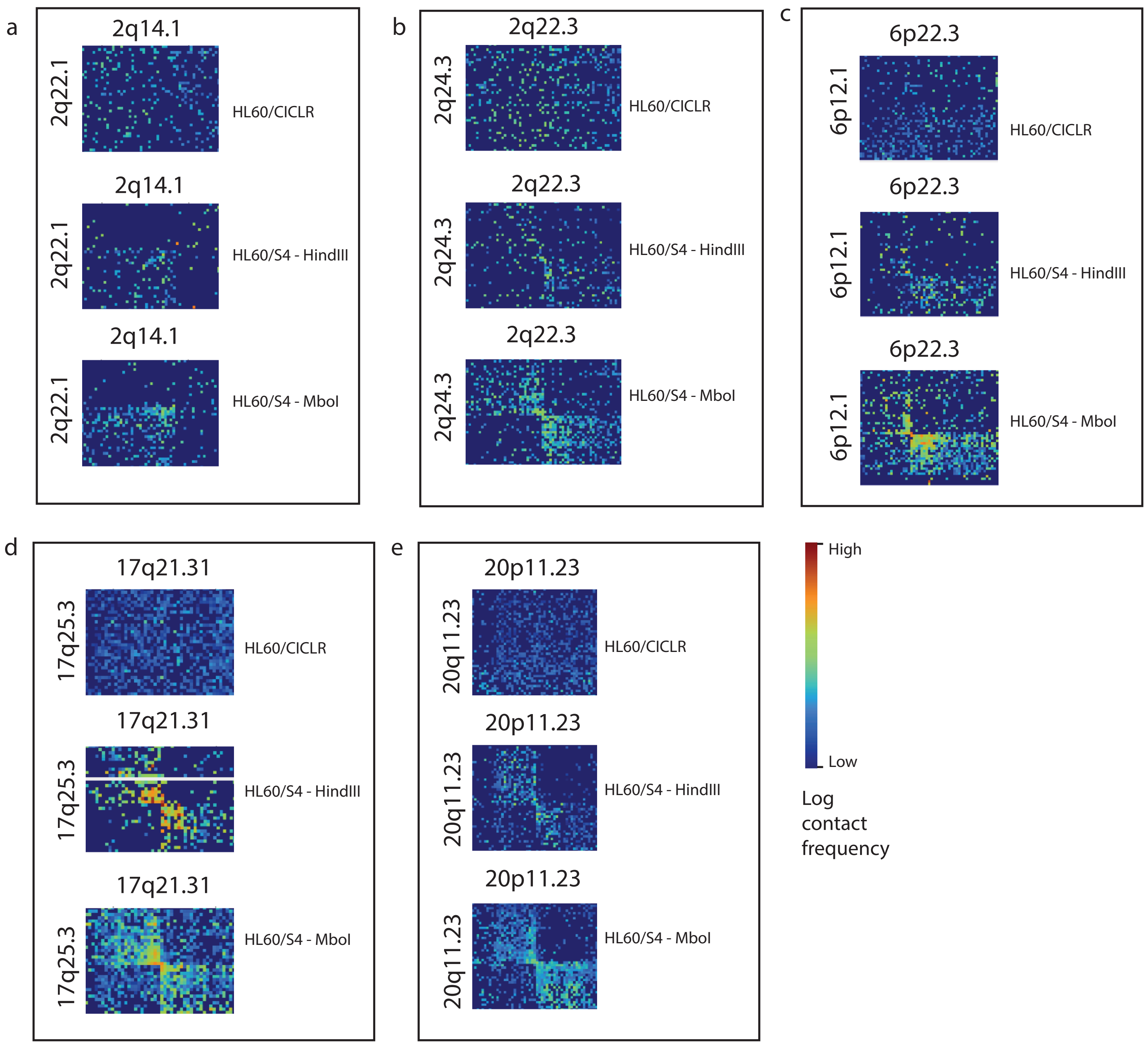

### Supplemental figure 5

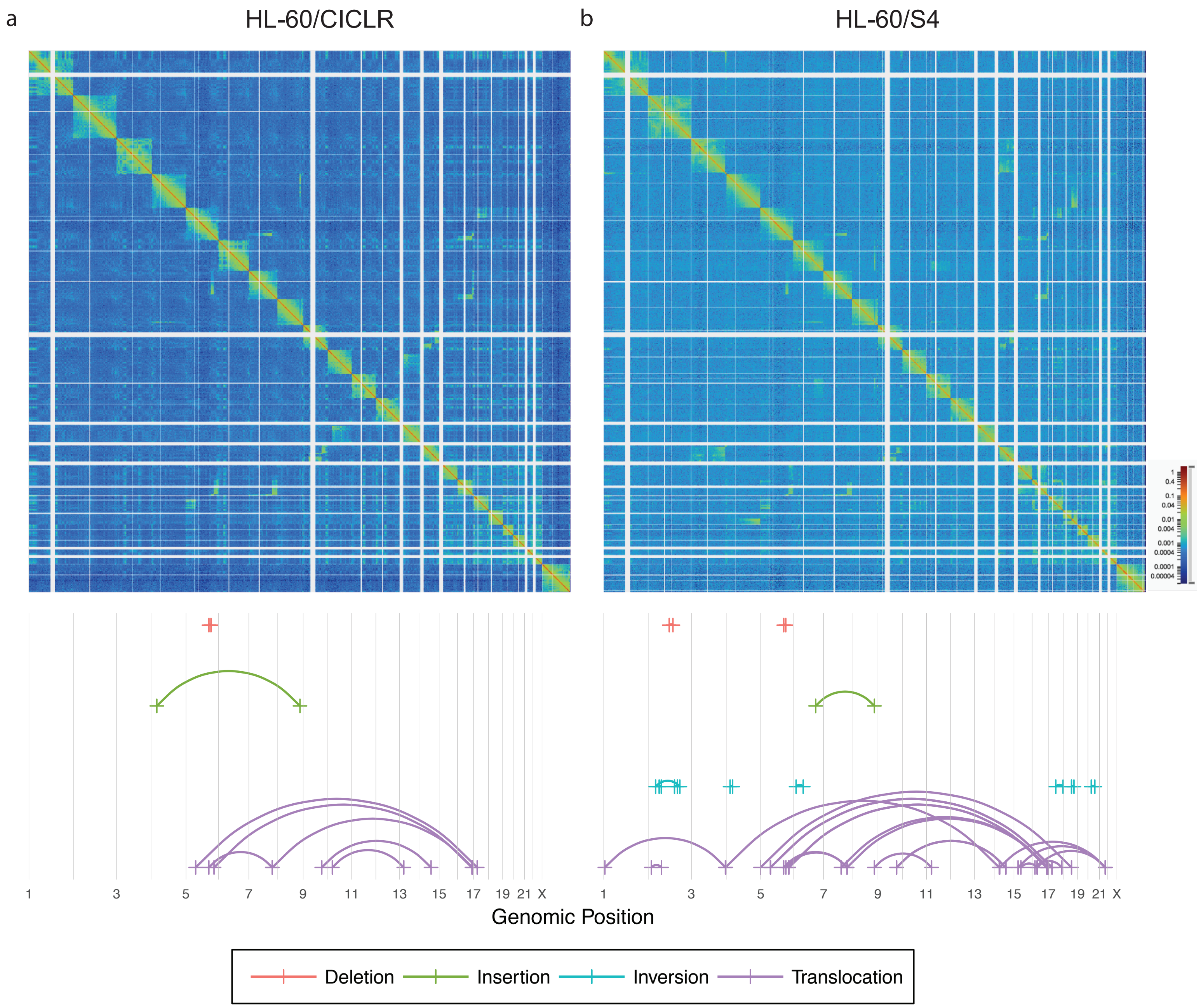
