## Supplemental methods for "Hi-C detects novel structural variants in HL-60 and HL-60/S4 cell lines"

### SUPPLEMENTARY METHODS

**Hi-C library preparation**

Cells were lysed in lysis buffer (500 μl 10 mM Tris-HCl pH 8.0, 10 mM NaCl, 0.2% NP-40 and 50 μl protease inhibitors, 15 min) on ice and disrupted using a Dounce homogenizer (10X, 1 min on ice, 10X). Lysed cells were pelleted (4°C, 5,000rpm, 3 min), washed twice and suspended in 1X restriction enzyme buffer, divided into five tubes (final volume of 362μl). SDS was added (1% FC), incubated (65°C, 10 min), placed on ice and quenched with Triton X-100 (10% FC). Chromatin was digested with HindIII (400U, 37^O^C, 16 h with rotation).

Restriction site overhangs were filled with biotin labeled Cytosine (1.5 μl 10 mM dATP, 1.5 μl 10 mM dGTP, 1.5 μl 10 mM dTTP, 37.5 μl 0.4 mM biotin-14-dCTP, and 50U Klenow, 45 min, 37°C). Tubes were placed on ice and reactions inactivated by SDS (10%), incubated (65°C, 30 min), and placed on ice immediately. Biotinylated fragments were ligated (4 h, 16°C) following addition of 7.61 ml ligation mix (745 μl 10% Triton X-100, 745 μl 10x ligation buffer [500 mM Tris-HCl pH 7.5, 100 mM MgCl_2_, 100 mM DTT], 80 μl 10 mg/ml BSA, 80 μl 100 mM ATP and 5.96 ml water), T4 DNA ligase (50U). Crosslinks were reversed and protein degraded by treatment with Proteinase K (25 μl, 12 mg/ml) at 65°C (16 h). After 16 h, additional Proteinase K (25 μl 20 mg/ml) was added to samples before a further incubation (2 h, 65°C).

Reactions were cooled to RT and DNA was purified by phenol (1X) and phenol:chloroform (1:1, 1X) extraction. The aqueous phase was transferred to high-speed centrifugation tubes (Oakridge). The DNA was ethanol precipitated (2.5 volumes 100% Ethanol and 1/10^th^ volume 3M Sodium Acetate), mixed by inversion, chilled (>30 min, -80°C), centrifuged (20 min, 10,000g, 4°C) and the DNA pellet dissolved in 450 μl 1x TE (10 mM Tris-HCl pH 8.0, 1 mM EDTA). The DNA was phenol:chloroform (1:1) extracted twice more with centrifugation (5 min, 14,000rpm, RT). The DNA was ethanol precipitated (2.5 volumes 100% Ethanol and 1/10^th^ volume 3M Sodium Acetate), mixed by inversion, chilled (>30 min, -80°C) and centrifuged (20 min, 10,000g, 4°C). The DNA pellet was washed with ice-cold 70% ethanol, centrifuged (10 min, >10,000g, 4°C), dried (~10 min) and suspended in 25 μl 1x TE. RNA was degraded by the addition of RNase A (1 μl 1 mg/ml), incubation (30 min, 37°C) before pooling the Hi-C libraries.

Biotin was removed from unligated ends using T4 DNA polymerase. ~5μg of Hi-C library was mixed with T4 DNA polymerase reaction buffer (1 μl 10 mg/ml BSA, 10 μl 10x NEBuffer 2, 1 μl 10mM dATP, 1 μl 10mM dGTP) and T4 DNA polymerase (5U) in a total volume of 100 μl before incubation (2 h, 12°C). Multiple 5 μg reactions were performed. Reactions were stopped by the addition of EDTA (2 μl 0.5M EDTA pH 8.0), reactions pooled and the DNA purified by three phenol pH 8.0:chloroform (1:1) extractions with centrifugation (5 min, 14,000 rpm, RT). DNA was Ethanol precipitated (2.5 volumes 100% Ethanol and 1/10^th^ volume 3M Sodium Acetate), mixed by inversion, chilled (>30 min, -80°C) and centrifuged (20 min, 10,000g, 4°C). The DNA pellet was washed with 70% Ethanol, dried, and suspended in a total volume of 100 μl H_2_O.

The DNA was sheared to a size of 300-500bp using a Covaris S2 instrument (duty cycle 5, intensity 5, cycles/burst 200, time 60 secs for 4 cycles). The sheared DNA ends were repaired (14 μl 10x ligation buffer, 14 μl 2.5 mM dNTP mix, 5 μl T4 DNA polymerase, 5 μl T4 polynucleotide kinase, 1 μl Klenow DNA polymerase and 1μl H_2_O; 30 min, RT).

DNA was purified using a Qiagen MinElute column according to the manufacturer's recommendations and the DNA eluted twice with Tris-low-EDTA (TLE: 15 μl 1x [10 mM Tris pH 8.0, 0.1 mM EDTA]). An ‘A’ tail was added to the DNA fragments using Klenow (5 μl [10x NEBuffer2, 10 μl 1 mM dATP, 2 μl H_2_O and 3 μl Klenow (exo-)]; 30 min, 37°C). The Klenow was inactivated (20 min, 65°C) and the reaction cooled on ice.

Reaction volumes were reduced to 20 μl using a speedvac, the DNA loaded on a 1.5% agarose gel with 1X TAE, run for 3 hours at 80V and DNA fragments between 300 and 500 bp were excised. DNA was purified using a Qiagen gel extraction kit according to the manufacturer’s recommendations and the DNA eluted with 50 μl 1x TLE. Purified DNA was combined in a final volume of 300 μl 1x TLE and the DNA concentration determined by QuBit.

To enrich for biotin-labeled fragments magnetic streptavidin beads (100 μl) were washed twice with Tween Buffer (TB: 400 μl [5 mM Tris-HCl pH 8.0, 0.5 mM EDTA, 1 M NaCl, 0.05% Tween]) and suspended in a volume of 2x No Tween Buffer (10 mM Tris-HCl pH 8.0, 1 mM EDTA, 2 M NaCl) equivalent to the remaining Hi-C library volume. All subsequent steps were performed in DNA LoBind tubes. Streptavidin beads were combined with Hi-C DNA and incubated (15 min, RT, slow rotation). Tubes were placed on a magnetic particle concentrator, the supernatant removed, the beads washed in NTB (NTB: 400 μl [5 mM Tris-HCl pH 8.0, 0.5 mM EDTA, 1 M NaCl]) and then with ligation buffer (100 μl 1x). The beads were suspended in ligation buffer (30 μl 1x) and transferred to a new tube.

The DNA bound to the streptavidin beads (~100 ng) was used for sequencing library preparation (Trueseq nano DNA LT kit, Illumina). Resuspension buffer (2.5 µl), DNA adaptor index (2.5 µl) and Ligation Mix 2 (2.5 µl) were added to the tube, and incubated (30°C, 2 h). Stop Ligation buffer (5 µl) was added to the tube to stop the reaction. Non-ligated adapters were removed by placing the tube on a magnetic particle concentrator and washing the beads twice with TB (400 μl 1x), once with NTB (200 μl 1x) and twice with NEBuffer 2 (200 μl and 50 μl 1x). The beads were suspended in 50 μl 1x NEBuffer 2 and transfered to a new tube.

The number of PCR cycles necessary to generate enough product for sequencing was determined by setting up four PCR reactions with 6, 9, 12 or 15 cycles (9 µl of sample [8 µl 1x NEBuffer2 + 1 µl DNA bound beads], 9 µl Enhancer PCR mix and 2 µl PCR cocktail. PCR conditions were: 95°C – 3 min, {98°C – 20 sec, 60°C – 15 sec, 6-15 cycles 72°C – 30 sec} 6-15 cycles, 72°C – 5 min, 4°C –hold. The optimal cycle number was determined by running the PCR reactions on a 5% polyacrylamide gel (60 min, 80 V). The remainder of the Hi-C-library-bound streptavidin beads were amplified in a large-scale PCR with the optimal number of PCR cycles. PCR products from the separate wells were pooled and the beads reclaimed. 1% of the large-scale PCR product was kept separate to run on a gel. The remainder of the PCR product was purified with Ampure beads (1.8x volume) according to the manufacturer's recommendations. DNA was eluted with TLE (50 μl 1x) and the Ampure bead purified PCR product compared to the original PCR product on a 5% polyacrylamide gel to insure the successful removal of the PCR primers. The Hi-C libraries were sequenced by Illumina paired end sequencing.

**Quality Control of Hi-C Data**

Biases due to restriction enzyme cutting, GC content, and sequence uniqueness can result in artifacts in the Hi-C libraries [1]. It is possible to handle these artifacts *in silico* by filtering out the offending reads. For example, one common artifact that arises is the insert size being above or below the expected size, as defined by the sonication during the library preparation. In this case, these reads are removed.

If the reads are of the appropriate size, then HiCUP [2] will check for several additional artifacts that can arise from the Hi-C protocol. For example, when the paired reads do not map to adjacent genomic restriction fragments, the read is filtered. If the paired-end reads map to the same restriction fragment the fragments have circularized, ligating to themselves. These fragments will still form linear molecules possessing a valid Hi-C junction, but they are not true evidence of a 3D spatial connection in the nucleus. Reads can also be identified as read pairs that map to the same genomic restriction fragment and are orientated away from or towards each other. If the DNA fragment end overlaps a restriction fragment cut site, it is likely that there was a failure to remove biotin residues from non-ligated DNA fragments, thus these reads also need to be removed.

In our experiments, the only artifact of note was the internal fragment error in the Hi-C alignment step. It is likely that the Hi-C protocol could have been better optimized to prevent the production of these internal fragments, which likely resulted from DNA restriction digestion at non-canonical sites. However, this QC check only means that a proportion of the reads were removed and does not affect the quality of the reads passed onto the analysis step. Following quality control, all five of the sequenced samples had sufficient valid reads (>30M valid read-pairs) for analysis.

Additionally, a valid Hi-C run should result in a certain proportion of reads that inform on *cis* versus *trans* connections, knowing that the Hi-C process favors proximity-dependent, intra-complex ligation, but also allows for exploration of longer distance trans-connections. HiCup best practices [2] recommends that the ideal experiment would have 55% of the aligned read-pairs representing inter-chromosomal interactions, 30% long-distance inter-chromosomal interactions (>20 kb apart), and 15% short-distance inter-chromosomal interactions (<20 kb apart). In our work, this distribution was slightly favoring inter-chromosomal interactions, representing 69% of the interactions in our experiment.
